# Signatures of Microbial Diversity at Multiple Scales of Resolution within Engineered Enrichment Communities

**DOI:** 10.1101/2022.10.01.510452

**Authors:** Elizabeth A McDaniel, Francisco Moya, Diana Mendez, Coty Weathersby, Ben O Oyserman, Jason Flowers, Shaomei He, Francesca Petriglieri, Caitlin Singleton, Per H Nielsen, Katherine D McMahon

**Author notes:** corresponding authors, email addresses and.

## Abstract

Microbial community dynamics are dictated by both abiotic environmental conditions and biotic interactions. These communities consist of individual microorganisms across the continuum of phylogenetic diversity, ranging from coexisting members of different domains of life and phyla to multiple strains with only a handful of single nucleotide variants. Ecological forces act on a shifting template of population-level diversity that is shaped by evolutionary processes. However, understanding the ecological and evolutionary forces contributing to microbial community interactions and overall ecosystem function is difficult to interrogate for complex, naturally occurring microbial communities. Here, we use two time series of lab-scale engineered enrichment microbial communities simulating phosphorus removal to explore signatures of microbial diversity at multiple phylogenetic scales. We characterized microbial community dynamics and diversity over the course of reactor start-up and long-term dynamics including periods of eubiosis and dysbiosis as informed by the intended ecosystem function of phosphorus removal. We then compared these signatures to lineages from full-scale WWTPs performing phosphorus removal. We found that enriched lineages in lab-scale bioreactors harbor less intra-population diversity than lineages from the full-scale WWTP overall. Our work establishes a foundation for using engineered enrichment microbial communities as a semi-complex model system for addressing the fundamental ecological and evolutionary processes necessary for developing stable microbiome based biotechnologies.

## INTRODUCTION

The abundance dynamics of individual taxa within a larger community are driven by many different forces, such as environmental fluctuations, competition for shared resources, cooperation, and/or predation events (1). Additionally, these ecological drivers act within the context of concurrent evolutionary changes because microbial ecological forces operate on similar time scales (2–4). A grand challenge in microbial ecology is understanding how diversity is structured within communities and how population dynamics change over spatiotemporal scales. Understanding the interplay of ecological and evolutionary forces that shape microbial communities is crucial for rationally designing and controlling diverse microbiomes for desired outcomes.

Defining the scope of microbial diversity in complex communities is currently limited by the technological constraints of cultivation-independent approaches. 16S rRNA gene amplicon sequencing surveys are currently the most commonly applied method to characterize the composition and diversity of microbial communities (5, 6). However, even full-length 16S rRNA gene sequencing cannot resolve populations at the species level for all taxa (7–9), and nearly identical 16S rRNA gene sequences can correspond to vastly different levels of divergence at the genome level (10–13). Genome-resolved metagenomic approaches have now been applied to virtually every habitat on earth to reconstruct population genomes of microbial lineages, providing both taxonomic affiliation and functional repertoire (14–16). Recruiting metagenomic reads back to assembled genomes has demonstrated that bacterial populations are organized as discrete genetic clusters referred to as “sequence-discrete populations” (SDPs), which are defined by a genetic discontinuity of a population in relation to the rest of the community (17–19).

Evidence for SDPs has been complimented by large-scale analyses of pairwise genome-wide average nucleotide identity (ANI) revealing a genome-based species threshold between 95-96% ANI (20, 21). Therefore, population genomes reconstructed from a mixed community can be designated as “species representatives” if two genomes are similar by less than 95% ANI. Furthermore, mapping metagenomic reads to sets of genomes above the 95% identity threshold has been used to analyze signatures of intra-population diversity and make evolutionary inferences (19, 22–24). These approaches have provided powerful insights into the ecology and evolution of microbial communities and uncultivated lineages (25).

Interrogating ecological and evolutionary dynamics in naturally occurring microbial communities is challenging due to the underlying community complexity and few cultivated representatives from these systems. Existing model systems in microbial ecology usually consist of 10 or fewer species, ranging from simple monocultures and enrichments (26), to low complexity natural microbiomes (25, 27), and synthetic communities (28, 29). These model communities allow for tractable systems to provide mechanistic insights into microbial dynamics within the context of a host and/or fluctuating conditions, and to link these fluctuations with overall ecosystem function (30). However, these models likely do not reflect dynamics in more complex systems such as freshwater, marine, and soil microbial communities. Therefore, new model microbial ecosystems of intermediate complexity are necessary to bridge detailed experiments designed to dissect *in situ* dynamics.

Engineered bioreactor systems have been used for decades to simulate bioprocesses and reproducibly enrich for simplified microbial consortia from a natural or engineered ecosystem (31–34). These systems are generally used to enrich low abundance microorganisms from a complex, natural environment for downstream cultivation (35) or further exploration of metabolic activities, regulatory controls, and evolutionary histories (36). Additionally, enrichment communities are often used as a platform for developing and testing new molecular and multi-omics technologies such as genome assembly approaches (37–39). The design of bioreactor systems allows for homogenously distributed substrates, reliable operation for long periods of time, and relatively well controlled conditions such as pH, carbon sources, and dilution rates (which control growth rates). Although studies usually focus on the highly enriched microbial lineage(s), other organisms than those of specific interest, termed are usually present in these bioreactor systems (40, 41). Therefore, these engineered systems represent semi-complex microbial communities that may be studied under controlled conditions.

To establish an enrichment community as a potential model system in microbial ecology and evolution in biotechnologies, fundamental patterns of community dynamics and diversity must be characterized and linked to ecosystem function. Here, we explored microbial community dynamics at multiple scales of genetic resolution using lab-scale enrichment bioreactors simulating Enhanced Biological Phosphorus Removal (EBPR) as a semi-complex system. EBPR is an economically and environmentally significant biotechnology implemented in numerous wastewater treatment plants globally for removing excess phosphorus from receiving waters. This process depends on polyphosphate accumulating organisms (PAOs) that incorporate inorganic phosphorus into intracellular polyphosphate (42, 43). Therefore in these ecosystems, phosphorus removal and the dynamic cycling of phosphorus are indicators of eubiosis. In contrast, when phosphorus removal and cycling is degraded, this can be a sign of dysbiosis. Long-term bioreactor experiments provide a controlled and tractable system where key ecosystem functions may be monitored, providing perfect laboratory ecosystems to study how ecological and evolutionary process connect to overall ecosystem function.

We first analyzed two time series of lab-scale bioreactors simulating EBPR designed to enrich for the well-studied PAO ‘*Candidatus* Accumulibacter phosphatis’. Numerous multi-omics approaches have been used to explore the ecology, gene regulation, and metabolic flexibility of Accumulibacter enrichments (11, 40, 41, 44–48). Using our time series, we analyzed population dynamics at multiple scales of resolution, specifically focusing on signatures of intra-population diversity using metagenomic reads mapping back to species-resolved population genomes. We explored these dynamics over the initial start-up period and long-term operation of two lab-scale bioreactor enrichments in which perturbations and community fluctuations occurred in the latter, and relate these back to phosphorus removal. We then compared these findings to a metagenomic dataset of 23 full-scale Danish WWTPs, from which approximately 500 high-quality species-representative genomes were reconstructed. We hypothesized that enriched lineages in lab-scale bioreactors would contain less intra-population diversity than those from the full-scale WWTPs. Overall, we provide an overview of microbial community dynamics and diversity at multiple scales of resolution in engineered enrichment bioreactors and how they relate to ecosystem function.

## MATERIALS AND METHODS

### Abigail and R1 Bioreactor Operation

For each bioreactor set, sequencing batch reactors with a 2-L working volume were inoculated with activated sludge from the Nine Springs WWTP in Madison, WI, USA in June of 2005 (Reactors R1 and R2) and February of 2021 (Reactor A named after the scientist Abigail Salyers). The Nine Springs WWTP in Madison, WI was operated to achieve both biological phosphorus and nitrogen removal using a modified University of Cape Town process (44). Each reactor was designed to simulate EBPR by primarily feeding with acetate as the sole carbon source, and operated in four daily cycles of 6 hours, with a hydraulic residence time (HRT) of 12 hours, and a solids retention time (SRT), or a mean cell residence time of 4 days (40). The total anaerobic/aerobic cycle time was 6 hours with 140 minutes anaerobic contact (sparging with N_2_ gas) and 190 minutes aerobic contact (sparging with oxygen), followed by a 30-minute settling period. Reactors were operated in either duplicate or triplicate to use as backups in case of the loss of EBPR in a bioreactor. Soluble phosphorus was characterized using Hach P-test powder assays and measured at the end of the anaerobic and aerobic phases for each sampling day (Hach, Loveland, CO).

The Abigail bioreactor was enriched through stepwise increases in the acetate loading after achieving complete acetate uptake by the end of the anaerobic phase, as monitored by HPLC assays. Similarly, the mean cell residence time was gradually decreased over the startup phase of operation to achieve stable enrichment of *Ca*. Accumulibacter phosphatis and phosphorus removal activity. All samples were collected from the Abigail bioreactor due to this bioreactor maintaining stable operation throughout the study. Reactor metadata are provided within the Abigail Figshare project repository at https://figshare.com/articles/dataset/Phosphorus_Metadata/21151645.

For the R1/R2 reactors, we aimed to understand microbial community dynamics and putative interactions over a period of mostly stable EBPR operation. These reactors were continuously operated with occasional upsets from June of 2005 to August of 2013 (44, 46, 49). On January 28^th^, 2010, the R1 reactor lost EBPR performance and was reseeded with biomass from the R2 reactor. We consider this Day 0 for all time series analyses of the R1 reactor. Therefore, all analyzed samples are from the R1 reactor during the period from January 28^th^ 2010 to April 30^th^ 2013, encompassing nearly 1200 days of operation and 300 mean cell residence times (generations) or turnover periods. Reactor metadata and chemistry-based performance data are provided within the R1 Figshare project repository at https://figshare.com/articles/dataset/R1_Chemical_Data_-_Subset/21151597.

### Sample Collection, 16S rRNA Gene Amplicon Sequencing, and Accumulibacter Clade Quantification

Unless noted, hereafter all procedures were performed identically for the Abigail and R1 reactor samples. Biomass samples were collected by centrifuging 2-ml of mixed liquor at 8,000 *x g* for 2 minutes and frozen at −80ºC as a pellet. Samples selected for DNA extraction and subsequent sequencing for the Abigail and R1 reactors are listed in the sample extraction status sheets within the respective Figshare project repositories. DNA was extracted for all samples using a modified phenol:chloroform bead-beating extraction protocol (40). Purified genomic DNA from all samples was sent to the University of Wisconsin – Madison Biotechnology Center for amplicon sequencing. The V3-V4 region of the 16S rRNA gene was amplified using the 341f and 805r primers and sequenced on an Illumina MiSeq with 2×300 bp paired-end reads. All raw sequences were filtered and amplicon sequence variants (ASVs) were identified using the DADA2 R package (50), with taxonomy assigned using the GTDB v86 database (51). The phyloseq R package was used to calculate and plot Shannon’s alpha diversity (52). The ampvis2 R package was used to visualize relative abundance over the time series (53). Accumulibacter clades were quantified in both reactor sets using established clade-specific primers targeting the *ppk1* locus (49, 54, 55). For the Abigail reactor, reads from two sequenced metagenomes described below were mapped to the existing UW Accumulibacter reference genomes (56, 57). Mapping results were used to assess Accumulibacter clade content and further confirm with clade IA, IIA, and IC primers from select samples. For the R1 reactor, monthly samples were used for qPCR with clade IA and IIA primers, as these were the dominant clades in this system (40, 44, 46).

### Metagenomic Assembly and Genome Curation

Two samples collected on operation days 43 and 60 from the Abigail bioreactor were submitted for metagenomic sequencing at the QB3 Sequencing Center at the University of California Berkeley. Metagenomic sequencing of select samples from the R1 bioreactor was performed at both the Joint Genome Institute (JGI) through a JGI Community Science Proposal (Proposal ID 873) and the QB3 Sequencing Center, all described in the table provided at https://figshare.com/articles/dataset/R1_Metagenomes_Information/21151633. Metagenomes sequenced at the JGI were from samples collected from the R1 bioreactor and were only used to aid in genome binning for differential coverage information. Additional metagenomes sequenced at QB3 were used for both metagenomic binning and subsequent intrapopulation diversity analysis. This resulted in a total of 15 metagenomic samples used for genome assembly for the R1 time series.

Unless otherwise noted, metagenomic assembly and genome curation was performed identically but in separate workflows for the R1 and Abigail metagenomes. All metagenomic samples were filtered with fastp and individually assembled into contigs using SPAdes v3.9.0 with the metagenomic option (58). Additionally, the two metagenomic samples from the Abigail reactor were coassembled with SPAdes v3.9.0. Reads from all metagenomic samples were reciprocally mapped to all assemblies using BBMap as part of the BBtools suite v38.07 with a 95% identity cutoff (59). Metagenome assembled genomes (MAGs) were binned using MetaBAT2 v 2.12.1 informed by the differential coverage of all samples only using contigs large that 1000 bp (60). Draft bins were preliminarily assessed for completion and contamination using CheckM v1.1.2 (61) and dereplicated into a set of non-redundant, species-representative bins using dRep v2.4.2 (62). All bins were classified using the genome taxonomy database (GTDB) using the GTDB-tk v.0.3.2 and database release 89 (51, 63). Accumulibacter bins from the R1 metagenomes were not reassembled through these efforts, and existing clade IIA (UW1) and IA (UW3) genome assemblies from Garcia-Martin et al. and Flowers et al. were used, respectively (40, 44). Bins assembled from the Abigail bioreactor and their relative abundance in the two samples are described in Table 1. Bins assembled from the R1 bioreactor are described in Table 2. The relative abundance of each bin from the R1 bioreactor in the 10 metagenomes used for population dynamics analyses are described in Table 3.

### Metagenomic Mapping, SNV Calling, and Microdiversity Metrics

Mapping to the concatenated set of dereplicate reference genomes for each bioreactor set was performed with bowtie2 v2.3.5.1, and relative abundance calculations were made with these mapping files using coverM v0.4.0 with the relative_abundance method. Preliminary coverage and breadth calculations were performed with inStrain v1.5.3 using the quick_profile method, which implements coverM (22). Using the mapping files generated from bowtie2, intrapopulation diversity was explored using the inStrain pipeline (22). Briefly, inStrain implements read filtering to meet the following criteria 1) both reads within a pair must map within 1500 bp of each other to the same scaffold, 2) The combined reads must map to the reference with at least 96% identity, and 3) At least one of the read pairs must have a mapq score >1, inferring that this is the unique, best mapping hit for the read pair (22, 23). To meet these criteria, all unpaired reads are removed, and paired reads must be mapped in the proper orientation with an expected insert size. To use a particular genome in a specific sample for downstream analyses, a genome had to have at least 10X coverage and at least 90% breadth in that sample, where breadth is the proportion of the genome that is covered (with most genomes with sufficient coverage containing greater than 95% breadth in a sample). To use a specific gene or identified SNV in downstream analyses, that gene or position had to have a coverage of at least 10X.

Nucleotide diversity π was calculated according to Nei et al. (64) for average nucleotide diversity and gene-specific nucleotide diversity. To measure the impact of homologous recombination in a population using metagenomic data, linkage disequilibrium is measured between pairs of SNV locations present between sets of paired reads (23, 65, 66). *r*^*2*^ and D’ were calculated according to VanLiere et al. (67), where SNVs used for calculations must have at least 10X coverage in that sample and are calculated between 20 sets of paired reads.

### Population Diversity across 21 Danish Full-Scale Wastewater Treatment Plants

23 full-scale WWTPs operating as EBPR plants were sampled across Denmark to produce over 1,000 high-quality MAGs (68). We only analyzed Illumina shotgun metagenomes sampled in 2018 as these had the highest sequencing depth. We analyzed metagenomes from 21 of these WWTPs because metagenomes from the Bjergmarken and Kalundbog WWTPs did not have sufficient coverage to most MAGs as compared to the other 21 samples. We included high-quality species-representative genomes from this dataset (completeness > 90%, redundancy < 10%) that were present in a sample with at least 10X coverage and 90% breadth. This stringent cutoff was applied to determine if a genome could be considered “present” in a sample among the Danish WWTPs, as species-representative genomes were originally assembled from individual WWTP metagenome assemblies. All mapping and inStrain calculations were performed as described above. Average nucleotide diversity across samples was calculated for a species across all samples that the species was detected in with sufficient coverage and breadth. The average nucleotide diversity of species within different phyla were compared for the full-scale WWTPs and to the other reactor datasets, averaging the nucleotide diversity for that species across samples in the other datasets in the same fashion.

### Data and Code Availability

All raw 16S rRNA gene amplicon sequencing reads, raw metagenomic sequencing reads, and non-redundant metagenome-assembled genomes are available through the SRA and Genbank linked to separate Bioproject accessions for the Abigail and R1 bioreactor datasets. All data for the R1 bioreactor is available at accession number PRJNA881607 and all data for the Abigail bioreactor is available at accession number PRJNA881609. The additional five R1 metagenomes sequenced at the JGI are available on IMG at accession numbers 3300026282, 3300026287, 3300026288, 3300026289, and 3300026302. Analyses of the POB metagenomes and genomes were performed on publicly available data at BioProject accession PRJNA704939. Analyses of the Danish WWTPs metagenomes and genomes were performed on publicly available data at BioProject accession PRJNA629478. All code is available on Github and mainly described in the MicDiv repository at https://github.com/elizabethmcd/MicDiv, with separate repositories for the specific Abigail reactor analyses at https://github.com/elizabethmcd/Abigail_EBPR_SBR and R1 time series analyses at https://github.com/elizabethmcd/R1PopDynamics. Reactor chemical metadata, sample extraction sheets, qPCR results, and metagenome information are provided in Figshare project repositories for the Abigail bioreactor at https://figshare.com/projects/Abigail_EBPR_Bioreactor/149212 and the R1 bioreactor at https://figshare.com/projects/R1_EBPR_Bioreactor_Long-Term_Timeseries/149209.

## RESULTS

### Succession Dynamics during Startup of a Lab-Scale Enrichment Bioreactor

We inoculated the Abigail bioreactor with full-scale activated sludge in 2021 and operated it for approximately 2 months to enrich for Accumulibacter (Figure 1). Over the course of reactor start-up and stabilization, we collected samples for 16S rRNA gene amplicon sequencing, quantitative PCR (qPCR) of the Accumulibacter polyphosphate kinase (*ppk1*) locus, and metagenomic sequencing (Figure 1A). The reactor exhibited good EBPR performance throughout most of the operation with the exception of a brief loss of phosphorus removal on day 44 (Figure 1B). The reactor was mainly comprised of Accumulibacter after approximately 2 weeks of operation as assessed by 16S rRNA gene amplicon sequencing (Figure 1C). We then assessed the Accumulibacter clade composition on operation days 43, 60, and 61 by qPCR of the *ppk1* locus. Accumulibacter has traditionally been designated into two types and several clades based on *ppk1* locus sequence identity (49, 54, 69), as the 16S rRNA gene is nearly identical among the known breadth of Accumulibacter clades (11, 44, 49). Although the nomenclature of Accumulibacter was recently redefined into several different species (70), we maintain the *ppk1*-based nomenclature throughout for ease of comparison to prior studies. The bioreactor was mainly comprised of clades IA, IC, and IIA (Supplementary Figure 1), representative strains of which were all previously characterized by various multi-omics analyses (11, 40, 44, 46, 47). Assessing within-sample alpha diversity using Shannon’s index showed a steady decrease in diversity over the first month of operation, and then fluctuating diversity dynamics afterward (Supplementary Figure 2B).

**Figure 1.**
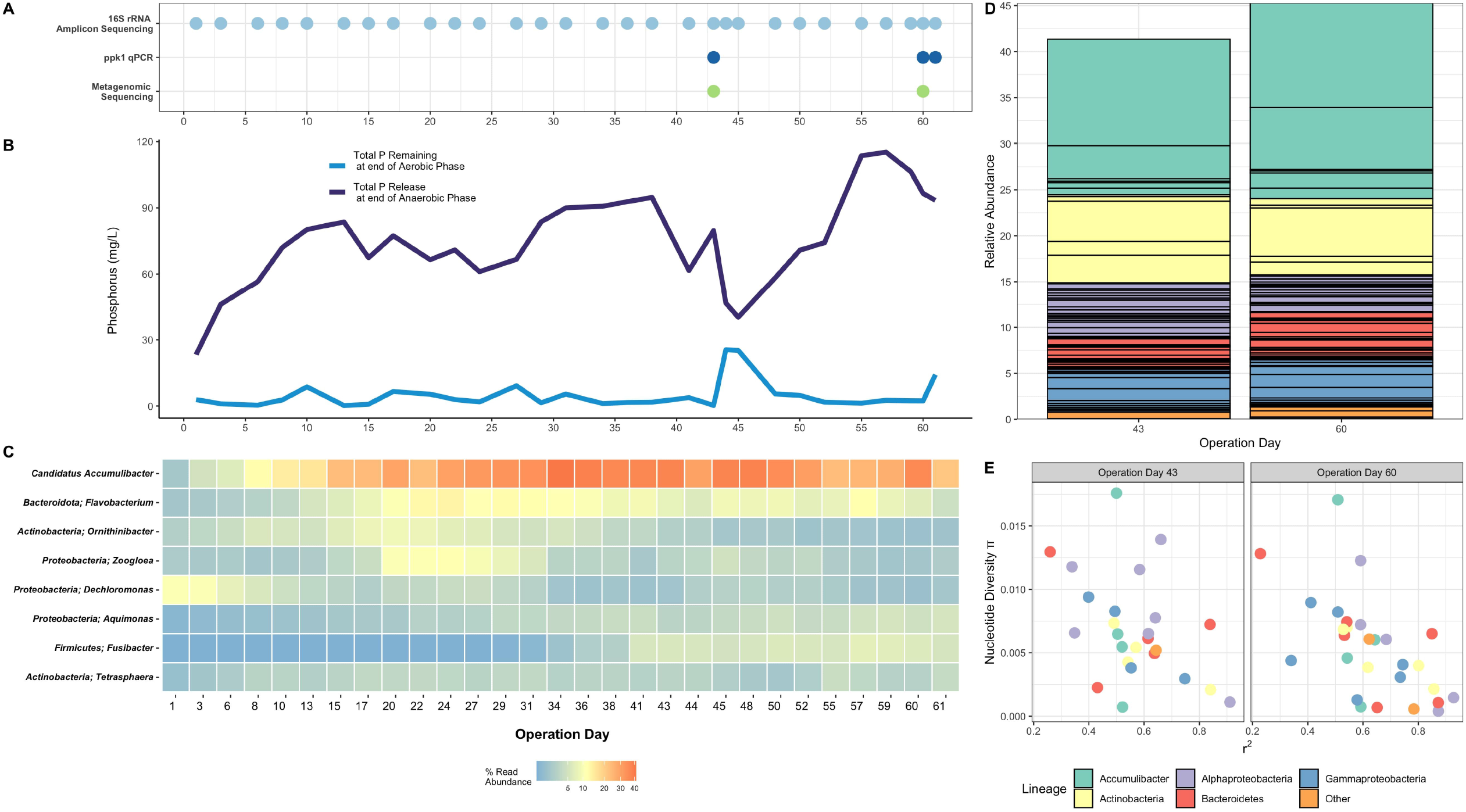
Succession Dynamics of an EBPR Lab-Scale Bioreactor (Abigail) at Start-up. **A)** Sampling scheme for 16S rRNA gene amplicon sequencing, qPCR of the Accumulibacter *ppk1* locus, and metagenomic sequencing. **B)** Phosphorus dynamics over the start-up phase, displaying both the total phosphorus release at the end of the anaerobic phase and total phosphorus remaining at the end of the aerobic phase. In EBPR-focused research, these dynamics are used as indicators of good or poor performance, where good constitutes high anaerobic-phase phosphorus release and low phosphorus remaining at the end of the aerobic phase. **C)** Population dynamics characterized by 16S rRNA gene amplicon ASVs at the genus level taken every few days. Color of each tile refers to the relative abundance of all ASVs within that genus. **D)** Relative abundance of MAGs assembled on operation days 43 and 60 assessed by mapping metagenomic reads back to dereplicated species-resolved MAGs. Each block in the stacked bar is an individual genome, where the color refers to the lineage of that genome, with colors matching E and the legend below. **E)** Within-sample nucleotide diversity π and mean linkage disequilbrium *r*^2^ of MAGs with sufficient coverage in samples taken on operation days 43 and 60.

We then selected operation days 43 and 60 for metagenomic sequencing to assess the strains that were enriched and their signatures of intra-population diversity (Figures 1D and 1E). We selected these time points because we wanted to compare population dynamics before and after the reactor performance decline on operation day 44 (Figure 1B). We were able to assemble 5 Accumulibacter MAGs, including clades IA and IIA based on ANI similarity to other reference Accumulibacter genomes. We were unable to assemble a high-quality genome of Accumulibacter clade IC and therefore mapped to an existing reference genome of this clade that was previously enriched in a lab-scale bioreactor also inoculated from the Nine Springs WWTP (47). The other three assembled Accumulibacter genomes did not show high similarity to existing reference genomes. Of the recovered Accumulibacter genomes, the clade IC genome comprised approximately 11% of the metagenomic reads on both operation days, with the next most abundant Accumulibacter strain comprising 6% of the metagenomic reads on day 60, and the remaining Accumulibacter genomes each comprising 1% or less of the metagenomic reads (Figure 1D). We also recovered genomes of other important EBPR lineages that are involved in phosphorus removal (71, 72). A *Tetrasphaera* genome that comprised approximately 5% of the metagenomic reads on both days, and a *Dechloromonas* genome that was present at low abundance on both days (Figure 1D, Table 1).

We then explored intra-population diversity of select genomes with sufficient coverage in both samples by calculating the nucleotide diversity metric π (Figure 1E). Enriched lineages displayed a wide range of intra-population diversity on both days from a maximum of approximately 0.018 to nearly clonal. The Accumulibacter clade IC UW-LDO genome contained the most intra-population diversity on both days, with other Accumulibacter genomes with sufficient coverage displaying lower nucleotide diversity values (Figure 1E). Homologous recombination can be assessed in metagenomic data by measuring the linkage disequilibrium *r*^*2*^ between pairs of SNV locations between sets of paired reads (22, 23). In populations with high amounts of recombination between pairs of linked SNVs, *r*^*2*^ is expected to decrease as the distance between pairs of SNVs increases, a pattern referred to as linkage decay (23). These lineages displayed a range of *r*^*2*^ values on both days, and most lineages with sufficient coverage to be detected for both operation days contained consistent *r*^*2*^ values (Figure 1E).

### Long-Term Dynamics of an EBPR Enrichment Bioreactor Community

We next analyzed a time series of an established enrichment bioreactor community simulating EBPR over a period of approximately 1200 days (Figure 2). Throughout the time series, we collected samples for 16S rRNA gene amplicon sequencing, qPCR of the Accumulibacter *ppk1* locus, and metagenomic sequencing at differing intervals (Figure 2A). Over this period, the bioreactor exhibited stable EBPR performance with typical periodic upsets characterized by a decrease in total phosphorus release at the end of the anaerobic phase and/or a decrease in the total phosphorus uptake by the end of the aerobic phase. Major operational upsets occurred on days 206, 208, 446, 539, 617, 956, and 1105 (Figure 2B). Most operational upsets were resolved within 3-4 subsequent days.

**Figure 2.**
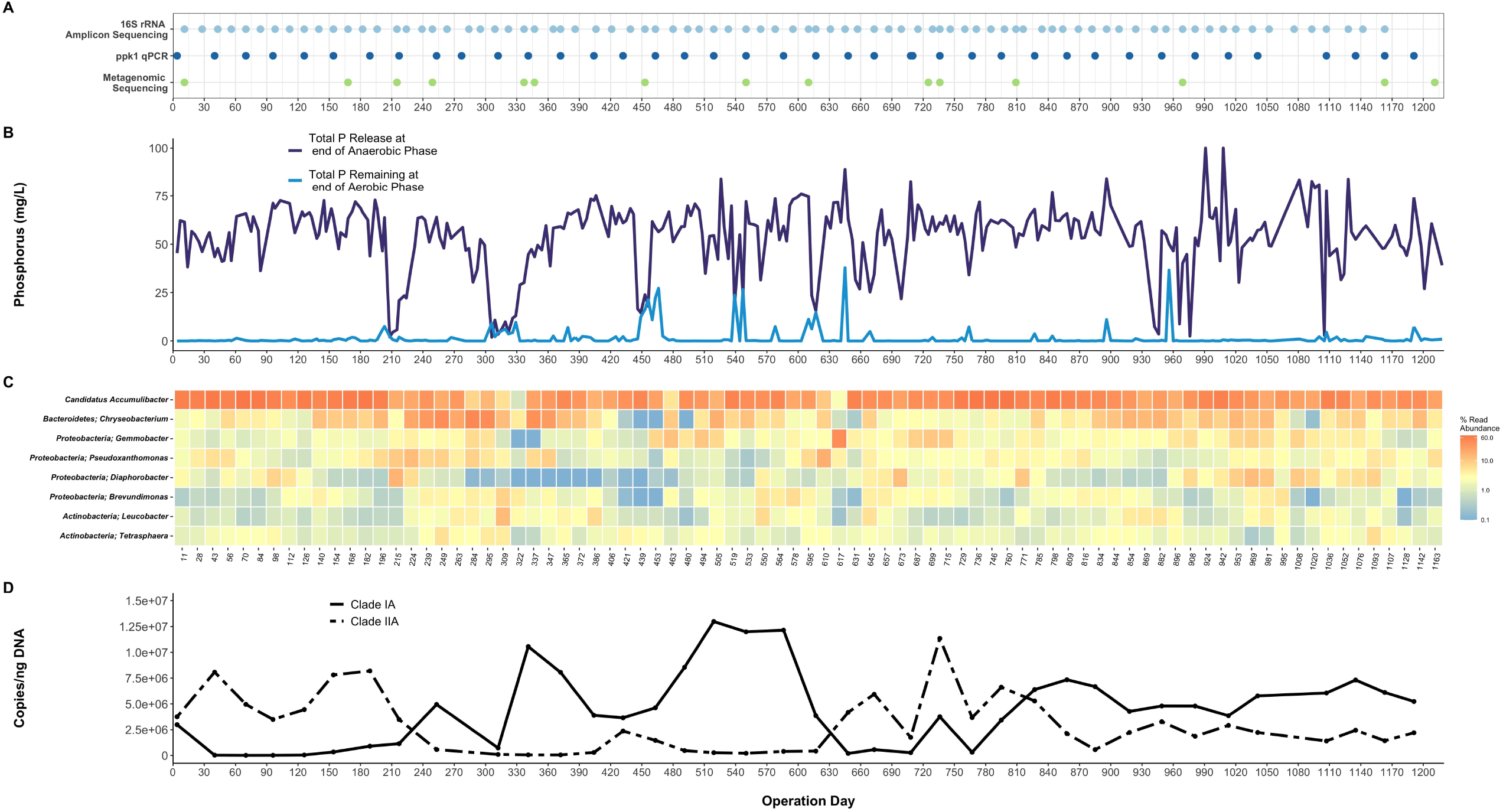
Long-Term Dynamics of a Lab-Scale Bioreactor (R1) Simulating EBPR Enriched with Accumulibacter. Operation days shown on the x axis for each panel are approximately spaced evenly for direct comparison between panels. **A)** Sampling scheme for 16S rRNA gene amplicon sequencing, qPCR of the Accumulibacter *ppk1* locus, and metagenomic sequencing. A total of 15 metagenomes were collected and all were used for *de novo* genome assembly. **B)** Phosphorus dynamics over bioreactor operation, displaying both the total phosphorus released at the end of the anaerobic phase and total phosphorus remaining at the end of the aerobic phase. Population dynamics assessed by 16S rRNA gene amplicon ASVs at the genus level from biweekly samples. Color of each tile refers to the relative abundance of all ASVs within that genus. Dynamics of Accumulibacter clades IA and IIA assessed by qPCR of the *ppk1* locus with clade-specific primers described by Camejo et al. (54).

### Community-Level Dynamics Assessed with 16S rRNA Gene Amplicon Sequencing

We first characterized coarse-level changes in the dynamics and diversity of the bioreactor community using 16S rRNA gene amplicon sequencing on biweekly samples (Figure 2C). Overall, Accumulibacter-ASVs dominated in most time points throughout operation, oftentimes comprising up to 50-60% of the reads (Figure 2C). Only for a few time points did Accumulibacter ASVs constitute less than 10% of the total read abundance. ASVs assigned to other abundant genera over the time series include the *Bacteroidetes* genus *Chryseobacterium*, the *Proteobacteria* genera *Gemmobacter, Pseudoxanthomonas, Diaphorobacter*, and *Brevunidmonas*, and the *Actinobacteria* genera *Leucobacter* and *Tetrasphaera* (Figure 2C). Previously using a trait-based comparative transcriptomics approach, we found that two *Pseudoxanthomonas* lineages exhibited similar *pstABCS* gene expression dynamics as Accumulibacter clade IIA (41), suggesting potential involvement in phosphorus uptake. Additionally, members of the Actinobacterial genus *Tetrasphaera* were experimentally verified as polyphosphate accumulating organisms and are abundant in Danish and global WWTPs (72–75). We then quantified alpha diversity within each sample using Shannon’s diversity index to survey how community diversity changed within samples over the time series (Supplementary Figure 2A). Decreases in alpha diversity preceded operational crashes on dates 208, 539, and 956 (Supplementary Figure 3A). Although there was a decrease in alpha diversity on operation day 617 in which Accumulibacter comprised less than 5% of the total relative abundance by 16S rRNA gene sequencing, there was a further decrease in alpha diversity that followed up until operation day 630 (Supplementary Figure 3A).

### Accumulibacter Dynamics Assessed with Clade-specific qPCR Primers

Using clade-specific *ppk1* qPCR primers we quantified the abundance of clades IA and IIA in monthly samples (Figure 2D). At the start of the time series at which this reactor was newly seeded from an existing enrichment bioreactor, clade IIA was more abundant. Although qPCR assays were only performed with monthly samples as compared to more frequent chemical sampling for phosphorus dynamics which occurred between every 3-7 days, a few clade fluctuations coincided with operational upsets for certain periods (Figure 2B and 2D). A flip from clade IIA to IA between operation days 189 and 217 coincides with the crash on operation day 208, and a flip from clade IA to IIA between operation dates 586 and 617 follows a series of operational upsets up to operation day 617 (Figure 2B and 2D). In contrast, a flip from clade IIA to IA between operation dates 827 and 858 does not appear to coincide with any obvious performance upset other than those that follow on operation days 956 and 1105 (Figure 2B and 2D).

### Snapshots of Population Dynamics using Metagenomic Sequencing

We then used 15 metagenomes sequenced from samples collected throughout the time series to assemble nonredundant, species-representative MAGs of microbial lineages within the system (Figure 2A, Table 2). We used the existing reference genomes for Accumulibacter clades IA (UW3-IA) and IIA (UW1-IIA) that were assembled previously from this bioreactor system (40, 44). These Accumulibacter genomes are no more than 85% similar by genome-wide ANI, and are therefore well below the sequence-based 95% ANI threshold for differentiating different species at the genomic level (20, 44). Additionally, clades IA and IIA of Accumulibacter were recently reevaluated and reclassified as separate species (70), but we retain the *ppk1*-based nomenclature throughout to aid in comparison to prior work. We assembled 167 bacterial species-representative MAGs of the bacterial flanking community (non-Accumulibacter) mostly spanning *Actinobacteria, Bacteroidetes*, and *Proteobacteria* phyla (Table 2). This includes several genomes of the experimentally verified and putative PAOs such as *Tetrasphaera, Ca*. Obscuribacter phosphatis, and *Dechloromonas*. Overall the assembled set of MAGs represent the majority of the community, as 80-90% of the metagenomic reads for each sample mapped back to the assembled genomes (Figure 3B). Of the 167 flanking community genomes recovered, 136 were more than 90% complete and less than 5% redundant as assessed with checkM. According to the MIMAG standard of a high-quality MAG requiring above 90% completion, less than 10% contamination, presence of all 3 rRNA genes (5S, 16S, and 23S) and at least 18 tRNA genes, 47 flanking bins met this criteria (76). Additionally, 55 bins met this completion and contamination requirement and had a full length 16S rRNA gene but were missing either of the two other rRNA genes.

**Figure 3.**
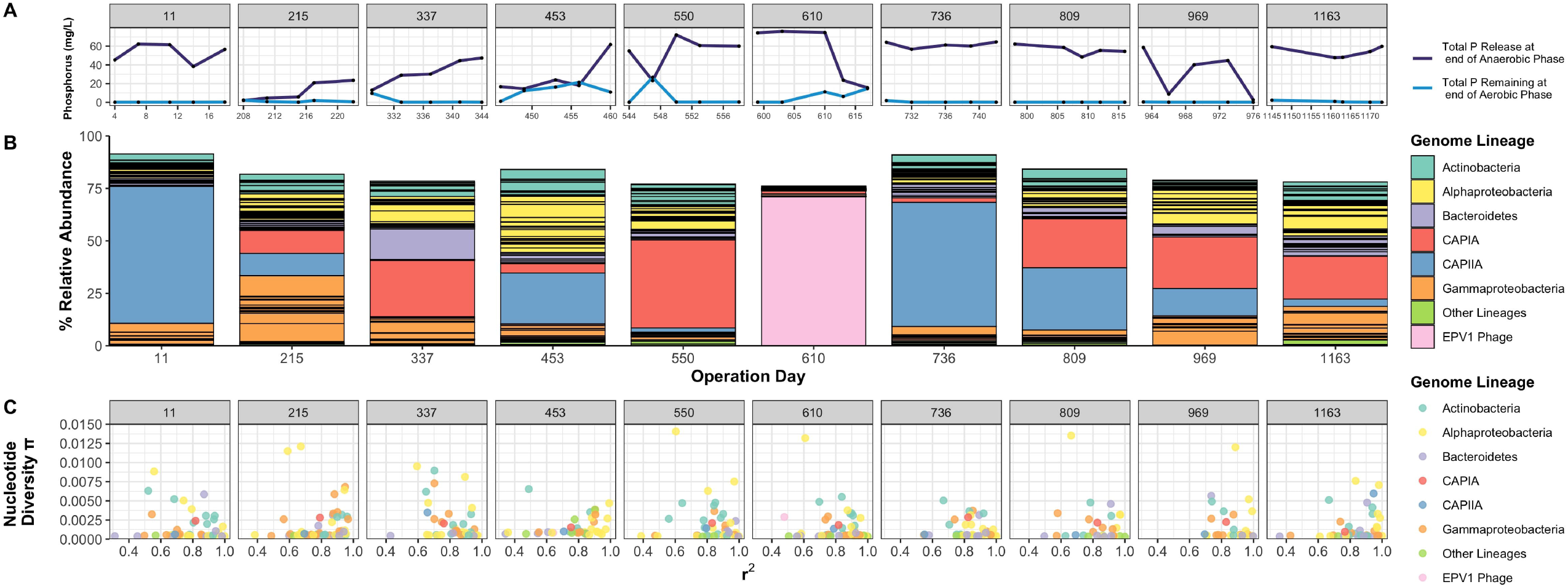
Snapshots of Population Diversity during Perturbations of a Lab-Scale Bioreactor Operated Long-Term. Metagenomes collected from the lab-scale bioreactor simulating EBPR operated long-term from operation days shown in Figure 3A. 10 specific metagenomes were chosen from samples where there was either a loss of EBPR performance or a recent flip in Accumulibacter clade abundance as assessed by qPCR of the *ppk1* locus in Figure 3D. **A)** Zoomed in phosphorus dynamics from Figure 3B showing phosphorus dynamics from the 2 days before and after the metagenome collection. Black points refer to the sample in which phosphorus dynamics were measured. **B)** Relative abundance of assembled MAGs assessed by mapping back metagenomic reads from each sample to the set of dereplicated species-resolved MAGs. Each block in the stacked bar refers to the relative abundance of an individual MAG, with the color corresponding to the lineage of that genome, with most lineages at the phylum level except for Accumulibacter and the EPV1 phage. Reads were mapped to the existing UW1-IIA (CAPIIA) and UW3-IA (CAPIA) genomes. **C)** Within-population diversity assessed by nucleotide diversity π and mean linkage disequilibrium *r*^2^ for MAGs with sufficient coverage and breadth in that sample. Colors of each point refer to the lineage of that genome, with most lineages at the phyla level except for Accumulibacter and the EPV1 phage. Colors for panels B and C match.

We specifically selected 10 metagenomes for sequencing based on Accumulibacter clade dynamics assessed through qPCR of the *ppk1* locus (Figure 2A). We were particularly interested in profiling the abundance and intra-population diversity of Accumulibacter and flanking community members around periods of operational upsets. Therefore, we selected timepoints either before, during, or after loss of EBPR performance coinciding with an Accumulibacter clade flip for metagenome sequencing (Figure 3). Samples collected on operation days 215, 337, 453, 550, 610, and 969 are characterized by a loss of EBPR performance and either exhibited a flip in Accumulibacter clade abundance or a crash and subsequent rise of that same clade (Figure 3). We mapped the metagenomic reads back from each of the 10 samples to the concatenated set of assembled genomes to assess the relative abundance of each population (Figure 4B). Towards the beginning of the time series on day 11 after seeding from a replicate enrichment bioreactor, Accumulibacter clade IIA constituted approximately 65% of the total reads mapped, with Accumulibacter clade IA comprising less than 1% of the total reads mapped. Clade IIA persisted in dominance throughout the first 200 days of operation.

**Figure 4.**
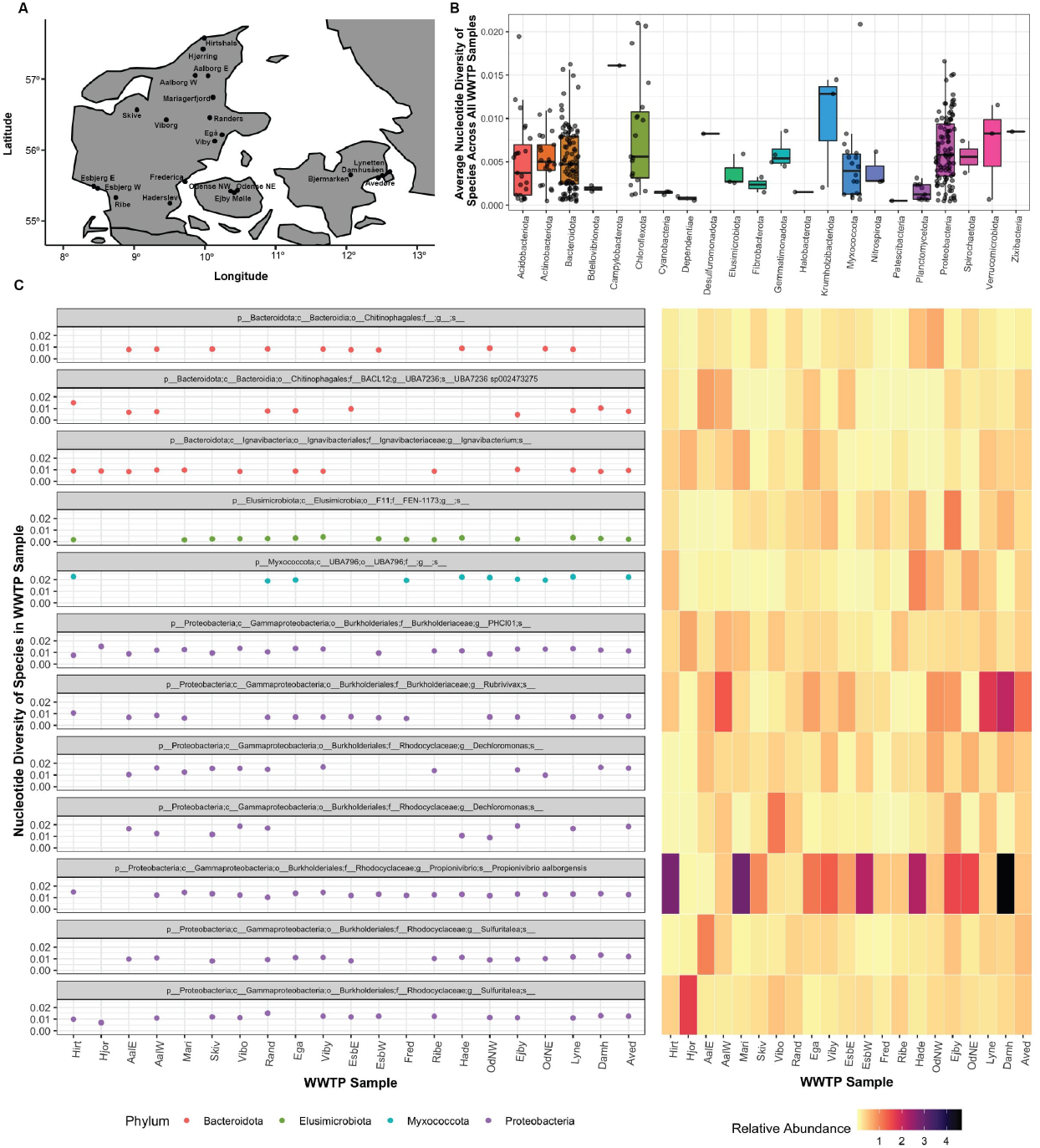
Population Diversity Across Danish Full-Scale WWTPs. **A)** Map of Denmark showing the location of each WWTP. Panels C and D are ordered on the x axis in the relative location of each WWTP, starting from the top down and going to the right, and are abbreviated with the first four letters of each WWTP location. **B)** Average nucleotide diversity of species across the full-scale WWTPs. For each species that had sufficient coverage and breadth in a sample, the nucleotide diversity of the species across all those samples was averaged and then grouped by the phyla of that species. Each point for each box plot is the average nucleotide diversity of an individual species in that phyla. **C)** Nucleotide diversity and relative abundance of the top 12 abundant lineages across the Danish WWTPs. Each row is an individual species, with the facet label on the left-hand side referring to the full taxonomy of that species. The left-hand side panel shows the within-sample nucleotide diversity of that species if there was sufficient coverage and depth of that genome in that sample, and the right-hand panel shows the relative abundance of that genome in that sample.

On day 208, EBPR performance was lost as evidenced by low anaerobic phosphate release (Figure 2B). On day 215, both Accumulibacter clades IA and IIA constituted 10% of the total reads mapped, respectively. Additionally, a *Pseudoxanthomonas* lineage within the *Gammaproteobacteria* constituted approximately 10% of the total reads mapped, followed by an *Alicycliphilus* lineage constituting 8.5% of mapped reads, a *Thauera* lineage constituting approximately 5% of mapped reads (Figure 3B). After this timepoint, the Accumulibacter abundance flipped from IIA to IA (Figure 2D). On day 307 there was a loss of EBPR performance characterized by a loss of phosphate release, and the abundance of clade IA decreased but did not flip to IIA. On day 337, Accumulibacter clade IA constituted approximately 26% of mapped reads, with a *Chryseobacterium* lineage constituting 14% of mapped reads (Figure 3B).

On day 403, there was a loss of EBPR performance characterized by both a loss of phosphate release and uptake. Based on the monthly *ppk1* qPCR assays, there was marked decrease in clade IA abundance, but Accumulibacter did not entirely flip to clade IIA. However on day 453, metagenomic reads mapped back to the MAGs resulted in 24% of the metagenomic reads mapping back to the IIA genome, and approximately only 4% to IA, indicating a clade flip around this time. However, clade IIA did not persist, as clade IA rose in abundance after this crash as seen in the *ppk1* qPCR data (Figure 2D), and 42% of the metagenomic reads mapped back to the clade IA genome on day 550 during a period of strong dominance (Figure 3B). Additionally on day 453, many individual genomes belonging to the *Alphaproteobacteria* rose in relative read abundance, including a *Beijerinkiaceae spp*. comprising 6% of mapped reads and a *Hyphomonadaceae spp*. comprising 3% of mapped reads. Two experimentally verified PAOs *Tetrasphaera* and *Dechloromonas* also rose in relative read abundance at this timepoint, comprising 4.5% and 3% of the metagenomic reads, respectively.

Around day 550, there was a transient loss of phosphate release and uptake. Afterwards there were two more upsets in EBPR performance, occurring on day 650 when the abundance of clade IA was previously stable, but dropped by day 650, upon which a rise in clade IIA followed (Figure 3D). From a metagenomic sample sequenced on day 610, 70% of the metagenomic reads mapped back to the EPV1 bacteriophage genome (Figure 3B). The EPV1 phage genome as well as two other complete phage genomes were previously recovered from these bioreactors several years earlier (77, 78). Within the EPV1 genome, a heat-stable nucleoid structuring (H-NS) protein was identified that was similar to a homolog in Accumulibacter, and was therefore hypothesized to be an Accumulibacter-specific phage (44, 77).

Following this crash, Accumulibacter clade IIA rose in abundance and constituted 60% of the metagenomic reads on operation day 736 (Figure 3B). Between days 800 and 850, there was a switch from Accumulibacter clade IIA to IA, but there was no obvious loss of EBPR performance, except for periodic moderate declines in total phosphate uptake (Figure 3B). The final two major operational upsets occurred at operation dates 956 and 1105, the former characterized by at first a loss of phosphate uptake followed by a loss of phosphate release, and the latter characterized by complete loss of phosphate release. During these upsets, there were no obvious fluctuations between clades IA and IIA, in which the relative abundance of the clade IA genome averaged between 20-25% of mapped reads at days 809, 969, and 1163, whereas the relative abundance of clade IIA decreased from 30% to 12% to 3.5% (Figure 3B).

We then profiled the *intra*-population diversity of Accumulibacter and flanking microbial community members. In each sample, we analyzed intra-population diversity for genomes that only had at least 10X coverage and 90% breadth of the genome (see Methods). Between a minimum of 30 to a maximum of 61 genomes met this threshold across the 10 samples, allowing for analysis of approximately 1/3 of the total genomes in most samples, including the EPV1 phage in two of the samples. Overall, most of the genomes did not contain high amounts of intra-population diversity as assessed by within-sample nucleotide diversity π (Figure 3C). In many samples, Accumulibacter clade IIA populations were not very diverse and were nearly clonal except for the last timepoint on operation day 1163. In contrast, the Accumulibacter clade IA population contained more intra-population diversity, averaging around 0.0025. Although both Accumulibacter clades IA and IIA reached a maximum relative abundance of approximately 45% and 65%, respectively, the nucleotide diversity of these populations in these timepoints was below 0.0025 (Supplementary Figure 4).

The most diverse populations over the time series belonged to the *Alphaproteobacteria* with the highest within-sample nucleotide diversity values on days 550, 610, and 809. A *Bosea* sp. contained within-sample nucleotide diversity values near 0.015 on days 550 and 610, and this was by far the most diverse population over the time series (Figure 3C). Another diverse *Alphaproteobacteria* population was a *Reyranella* sp. towards the beginning of the time series (Figure 3C). Notably, these populations were not very abundant compared to other lineages, comprising less than 1% of the total relative abundance in most samples, respectively Additionally, in two samples that the EPV1 phage was detected, the bacteriophage population was not that diverse across the genome, where the population was nearly clonal on operation date 453 and a within-sample nucleotide diversity value of less than 0.005 on operation date 610 when the phage bloom occurred (Figure 3B and 3C).

Homologous recombination can be assessed in metagenomic data by measuring the linkage disequilibrium *r*^*2*^ between pairs of SNV locations between sets of paired reads (22, 23). In populations with high amounts of recombination between pairs of linked SNVs, *r*^*2*^ is expected to decrease as the distance between pairs of SNVs increases, a pattern referred to as linkage decay (23). Additionally, the D’ statistic can also be measured as a function of recombination, where any value < 1 infers recombination occurring (67). Generally, the mean *r*^*2*^ and D’ values for a measured population follow a linear relationship (23, 67). This relationship was observed for several genomes that met the coverage thresholds in multiple timepoints, with most populations containing a mean *r*^*2*^ value close to 1 and an average D’ value at 1 (Supplementary Figure 5). In populations where recombination rates are low or if selection is strong, clonal strains can compete until one or more beneficial alleles are selected for, in which the single clonal genotype can sweep to fixation. This can purge diversity in a genome-wide sweep and results in a clonal expansion (79, 80).

### Population Diversity between Lineages in Full-Scale WWTPs and Enrichment Bioreactor Systems

*Intra-Population Diversity of Abundant Microbial Lineages in Danish Full-Scale WWTPs* To compare profiles of intra-population diversity in lab-scale lineages to those of full-scale lineages, we then analyzed a dataset of over 1000 high-quality genomes recovered from 21 Danish full-scale WWTPs (68) (Figure 4). We mapped back metagenomes from each WWTP sample collected from across Denmark in 2018 (Figure 4A) to the concatenated set of 581 high-quality, species-representative genomes and profiled the intra-population diversity across samples. We first surveyed the average genome-wide nucleotide diversity of species across all samples the genome was detected in above at least 10X coverage (Figure 4B). For some phyla, there were only a few or a single species representative present with enough coverage across all samples to profile within-sample nucleotide diversity, such as members of the *Bdellovibrionota, Cyanobacteria, Dependentiae, Desulfuromonadota, Halobacterota, Patescibacteria, and Zixibacteria* (Figure 4B). Other phyla contained several species with sufficient coverage in multiple samples and exhibited a range of intra-population diversity such as members of the *Acidobacteriota, Bacteroidota, Chloroflexota*, and *Proteobacteria* (Figure 4B).

We then examined the intra-population diversity of the most abundant lineages across the full-scale WWTPs (Figure 4C). This included 3 members of the *Bacteroidota*, one member of the *Elusimicrobiota*, one member of the *Myxococcota*, and 7 members of the *Proteobacteria*. Overall, intra-population diversity of a species was consistent across the full-scale WWTPs even when the relative abundance of that lineage varied in a WWTP, with some exceptions (Figure 4C). Among the most abundant lineages, a member of the *Elusimicrobiota* exhibited the lowest amount of nucleotide diversity across WWTPs, whereas a member of the *Myxococcota* exhibited the highest amount of nucleotide diversity across WWTPs (Figure 4C).

Several members of the *Rhodocyclaceae* within the *Proteobacteria* were detected in high abundance across WWTPs. Members within the *Dechloromonas* have been found to exhibit polyphosphate accumulating activity based on Raman-FISH experiments (81, 82). Two *Dechloromonas* species were detected in several WWTP samples at nearly 1% of the total reads, and exhibited varying within-sample nucleotide diversity. For example, a *Dechloromonas* species had a minimum of 0.01 nucleotide diversity in the OdNW and a maximum of nearly 0.02 nucleotide diversity in the Ejby and Vibo WWTPs. Taxa within the *Sulfuritalea* genus belong to the core microbial community in worldwide WWTPs and are involved in nitrogen transformations (83), and this species had more consistent nucleotide diversity patterns across WWTPs. The novel glycogen-accumulating organism ‘Ca. *Propionivibrio aalborgensis’* was found to be highly abundant in full-scale WWTPs, comprising approximately 3% of the total biovolume (84). Since glycogen-accumulating organisms do not store polyphosphate intracellularly, they compete with PAOs for similar carbon substrates without contributing to phosphorus removal. This lineage was one of the most abundant across all sampled WWTPs, and had more consistent nucleotide diversity patterns.

### Intra-Population Diversity across Different Types of EBPR Systems

We lastly compared patterns of intra-population diversity of microbial lineages within all three datasets surveyed as a part of this study (Figure 5). This includes metagenomes from the Abigail bioreactor sampled close to startup, the R1 long-term time series that was seeded from a prior enrichment, and the 21 Danish full-scale WWTPs. We also included a photosynthetically operated bioreactor (POB) simulating EBPR (57, 85) from which we reconstructed genomes of the bacterial community from three time-points near the end of bioreactor operation when stable EBPR performance was achieved (57). We recovered genomes of six different Accumulibacter clades, including clades IA, IIA, IIB, IIC, IID, and IIF (57). The reactor was mainly comprised of Accumulibacter and an *Ignavibacteria* lineage at these time points, with Accumulibacter clade IIC primarily enriched towards the end of operation.

**Figure 5.**
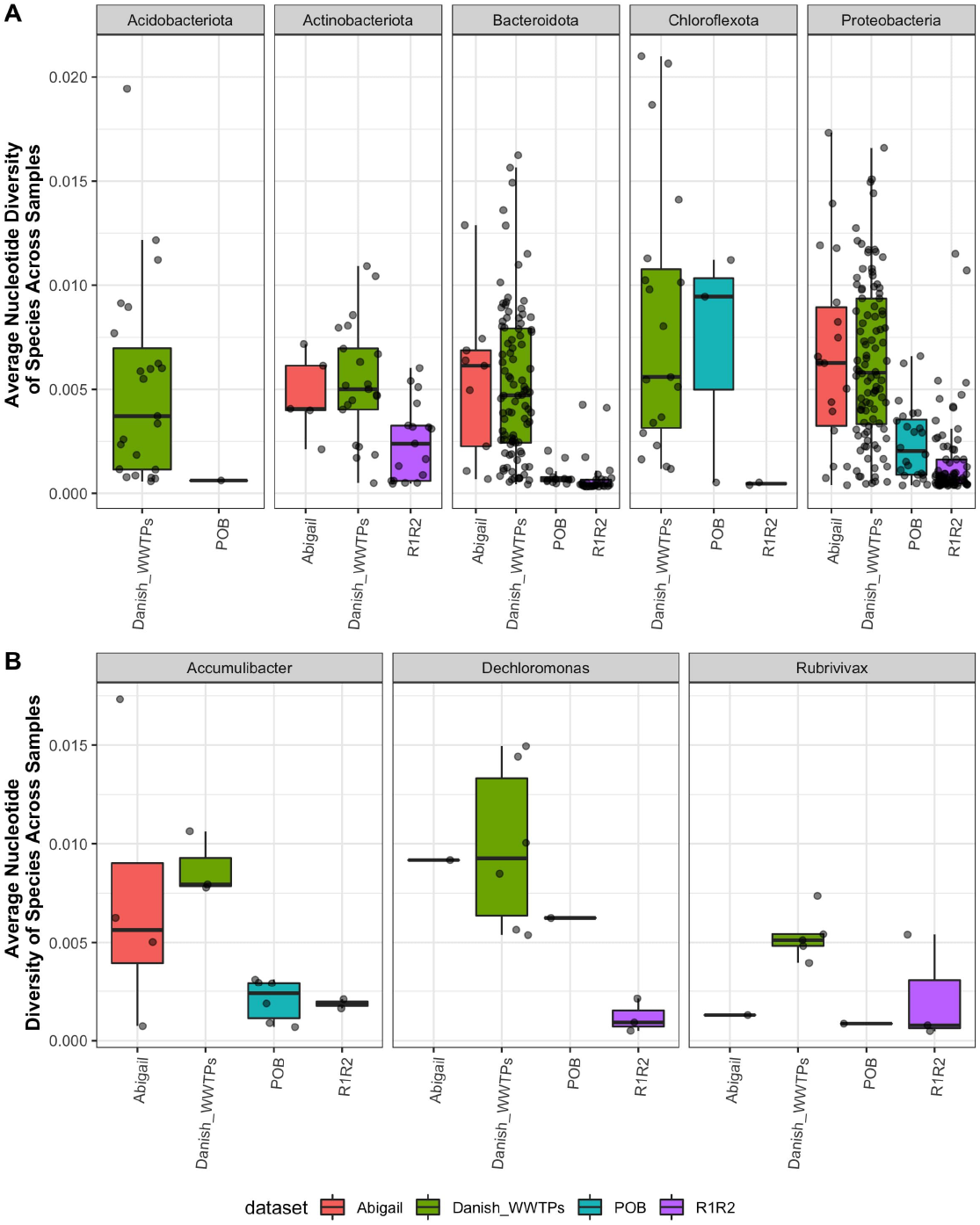
Comparisons of Population Diversity Across EBPR Systems. Comparing intra-population diversity of species surveyed in the Abigail, POB, and R1 lab-scale bioreactors and across the full-scale Danish WWTPs. As in Figure 5, for a species with sufficient coverage and depth in a sample, nucleotide diversity was calculated and then averaged for all other samples that met this criteria for the species. Average nucleotide diversity of the species is then shown within phyla, where the individual points are the average nucleotide diversity of that species. **A)** Average nucleotide diversity of species across EBPR systems shown at the phyla level. Species within the phyla Acidobacteria, Actinobacteria, Bacteroidota, Chloroflexota, and Proteobacteria were chosen as these phyla were represented the most across all four datasets. **B)** Average nucleotide diversity of species across EBPR systems shown at the genus level for genera that were in all four datasets. This included Accumulibacter, Dechloromonas, and Rubrivivax, whereas individual points for a dataset are individual species within that genus for that dataset.

We first compared average nucleotide diversity of species within the *Acidobacteriota, Actinobacteriota, Bacteroidota, Chloroflexota*, and *Proteobacteria* phyla (Figure 6A). The only phyla that contained lineages in all four datasets were the Bacteroidota and Proteobacteria. In support of our hypothesis that smaller, more selective systems harbor less diversity, lineages for phyla sampled across the Danish WWTPs contained the highest amounts of intra-population diversity, whereas the POB and R1 reactor datasets contained lower amounts of intra-population diversity (Figure 5A). This pattern is especially predominant when comparing lineages from these systems within the *Bacteroidota* and *Proteobacteria*. The Abigail system contained lineages with more variable levels of intra-population diversity, as might be expected as these samples were taken closer to start-up when the reactor was not fully enriched or operated for as long as the POB and R1 time series.

We then selected species of three genera that were present in all four datasets to compare their levels of intra-population diversity (Figure 5B). This included members of Accumulibacter, *Dechloromonas*, and *Rubrivivax*. The Abigail and R1 reactors were designed to enrich for Accumulibacter, and the POB reactor was seeded in part from a lab-scale bioreactor already enriched with Accumulibacter. For all three genera, the R1 dataset populations contained the least amount of intra-population diversity. Interestingly, the Abigail reactor had the most variable amounts of intra-population diversity for different clades or species of Accumulibacter, ranging from near clonal to above 0.015 (Figure 5B). For members within the *Dechloromonas*, we only recovered one species each from the Abigail and POB reactors, whereas there were several species of *Dechloromonas* across the Danish WWTPs with varying levels of intra-population diversity (Figure 5B). For species of *Rubrivivax*, the species from the Abigail and POB reactors contained very low amounts of within-population diversity, and the R1 reactor had three *Rubrivivax* species with varying levels of nucleotide diversity.

## DISCUSSION

In this work, we used two time series of lab-scale enrichment microbial communities simulating phosphorus removal to understand population dynamics in a semi-complex microbial community. By combining multiple cultivation-independent technologies such as 16S rRNA gene amplicon sequencing, qPCR of a focal lineage (*Ca*. Accumulibacter), and deep metagenomic sequencing, we were able to characterize signatures of microbial diversity at multiple scales of resolution in these enrichment communities. By comparing these signatures to lineages in full-scale WWTPs, we confirmed our hypothesis that enriched lineages in lab-scale bioreactors contain less intra-population diversity than that of full-scale lineages overall. Additionally, by exploring population dynamics during both the start-up phase of reactor operation and long-term dynamics of occasional operation upsets, we propose that lab-scale enrichment microbial communities simulating engineered processes may be an ideal model system for addressing fundamental questions at the interface of ecology and evolution.

Two grand challenges in microbial ecology include how microbial communities assemble in response to novel environments, and how community dynamics change over long periods of time. Examples of surveys of microbial community assembly and succession range from relatively simple communities such as those of cheese rinds (86) and pitcher plants (87), to complex ecosystems such as that of the infant gut microbiome (88) and full-scale WWTPs (89). Engineered enrichment communities provide appealing semi-complex systems for studies of community assembly and succession. Lab-scale bioreactors could be inoculated with complex seeds from activated sludge, anaerobic digesters, or other natural environments and modified with different carbon sources, mean cell residence time, and other parameters to test how communities are enriched at multiple scales of resolution. One shortcoming of our study is that we did not have corresponding samples from a full-scale WWTP and the enrichment bioreactor to directly characterize the succession dynamics and diversity signatures of the community. An interesting follow-up would be performing deep metagenomic sequencing from the complex inoculum through enrichment for tracking succession at multiple scales.

Additionally, our analysis of long-term dynamics of the R1 enrichment community demonstrate that lab-scale bioreactors could be used as semi-stable model systems for long-term studies of microbial community dynamics. Currently, most long-term time series of microbial communities are conducted with 16S rRNA gene amplicon sequencing due to both the costs of long-term sequencing studies and the underlying complexity of these communities (90, 91). Not only can lab-scale enrichment communities be maintained long-term, but the enrichment of a focal lineage in a semi-complex community could be an ideal system for understanding microbial interactions.

Engineered enrichment communities may also provide a foundation for developing new biotechnologies. Currently, synthetic communities are usually composed of 10 or fewer strains selected for specific functions (28, 29). To rationally engineer predictable and robust microbial biotechnologies, we must understand the driving forces of within-species variation, namely how this genetic variation is stratified within populations and is maintained or differs across spatiotemporal scales. In this work, we found that multiple closely related lineages such as different clades of *Ca*. Accumulibacter can coexist and fluctuate in abundance, most likely depending on how short the mean cell residence time is (92, 93). Although we found that most enriched lineages contain low amounts of intra-population diversity and are nearly clonal, this could be further explored by adjusting the carbon sources and mean cell residence time. The semi-complex nature of these enrichment communities could provide an ideal scenario for creating new predictable and robust biotechnologies by combining top-down and bottom-up approaches (94). Synthetic communities comprised of subsets of isolates from high-throughput cultivation experiments of the enrichment community could be paired with multi-omics experiments to bridge mechanistic experiments with *in situ* observations. The combination of top-down and bottom-up approaches for the same microbial ecosystem was recently applied to cheese communities to create a high-quality genome catalog and test succession patterns of subsets of community members (86). The additional appeal of using engineered enrichment communities is that the simulated process, such as phosphorus removal, denitrification, or methane production, could be used as an ecosystem parameter to screen subsets of communities or in response to perturbations.

## ACKNOWLEDGEMENTS

We thank the University of Wisconsin-Madison Biotechnology Center DNA Sequencing Facility for 16S rRNA gene amplicon library prep and sequencing. Metagenomic sequencing was provided in part through a Joint Genome Institute Community Science Proposal (Proposal ID 873). We thank the University of California Berkeley John Coates Sequencing Center for library prep and metagenomic sequencing. Special thanks to Phil Hugenholtz for early discussions that inspired this work. We thank the Villum Foundation (Dark Matter grant 13351) supporting the retrieval and analysis of MAGs from the Danish WWTPs. This work was supported by funding from the National Science Foundation (MCB-1518130) to KDM. Funding was provided to EAM by a fellowship through the Department of Bacteriology at the University of Wisconsin - Madison. This research was performed in part using the Wisconsin Energy Institute computing cluster, which is supported by the Great Lakes Bioenergy Research Center as part of the U.S. Department of Energy Office of Science (DE-SC0018409).

## TABLE LEGENDS

**Table 1. Abigail Bioreactor MAG Statistics.** Genome information including completeness, contamination, size, GC, GTDB taxonomy, and relative abundance in the two metagenomes for each genome. Includes mapping information for the UW-LDO genome that was not assembled as part of this study. Also available at https://figshare.com/articles/dataset/Table_1_-_Abigail_bins_table/21159001

**Table 2. R1 Bioreactor MAG Statistics.** Genome information including completeness, contamination, size, GC, GTDB taxonomy, number of rRNAs, tRNAs, and number of each type of rRNA for each genome. Also available at https://figshare.com/articles/dataset/Table_2_-_R1_bins_table/21159004

**Table 3. R1 Bioreactor MAG Relative Abundance.** Relative abundance of each genome in each of the 10 samples used for population dynamics and diversity analysis. Relative abundance was not analyzed for the 5 metagenomes for which differential coverage was only used to aid in assembly and binning and not for analyses. Available at https://figshare.com/articles/dataset/Table_3_-_R1_bins_relative_abundance_table/21159010

## SUPPLEMENTARY FIGURE LEGENDS

**Supplementary Figure 1.**
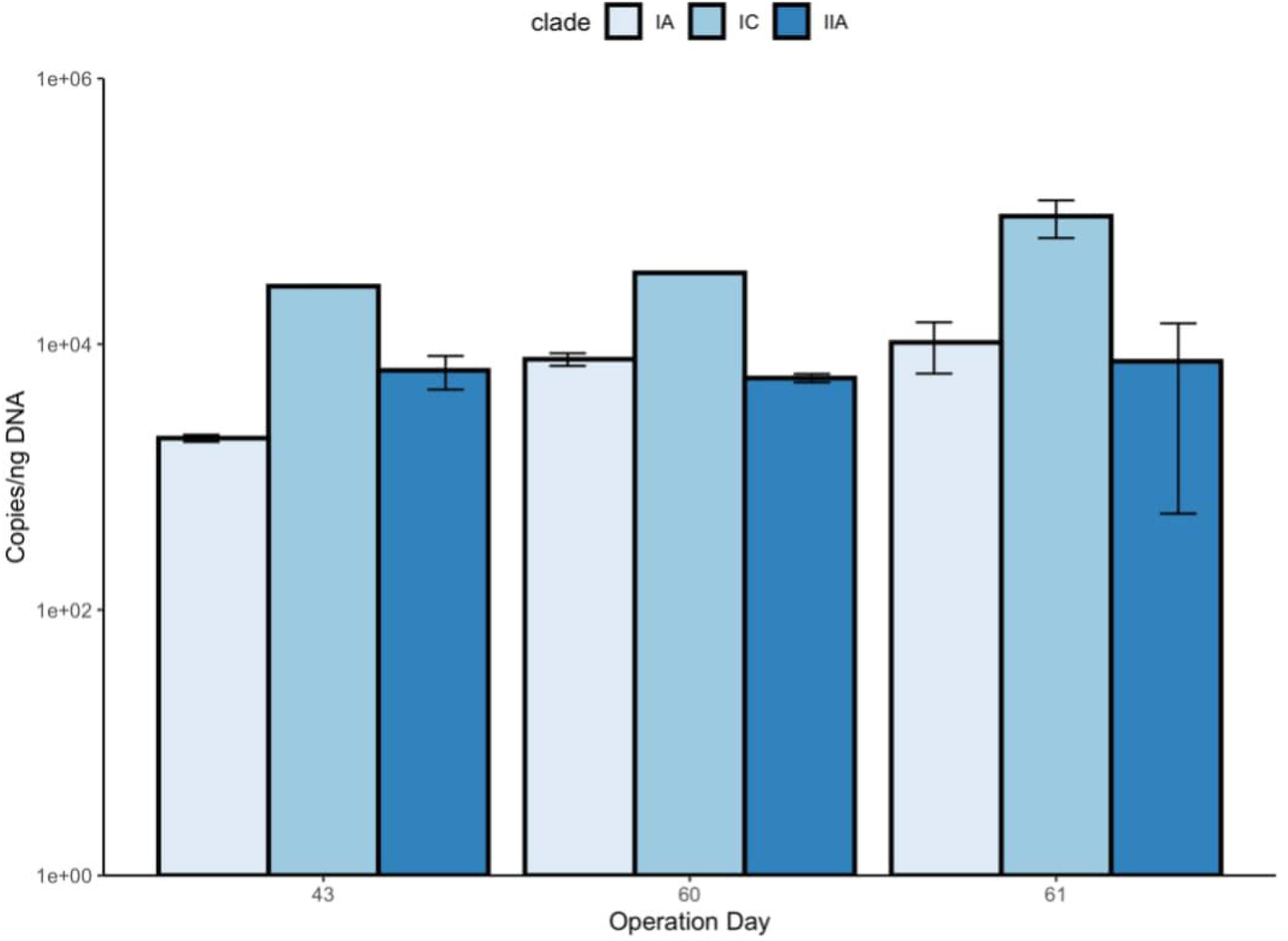
Quantifying Accumulibacter clades with clade-specific *ppk1* qPCR primers. Accumulibacter clades IA, IIA, and IC were quantified using clade-specific *ppk1* primers described in Camejo et al. (54). DNA was extracted from samples collected on days 43, 60, and 61 of bioreactor operation.

**Supplementary Figure 2.**
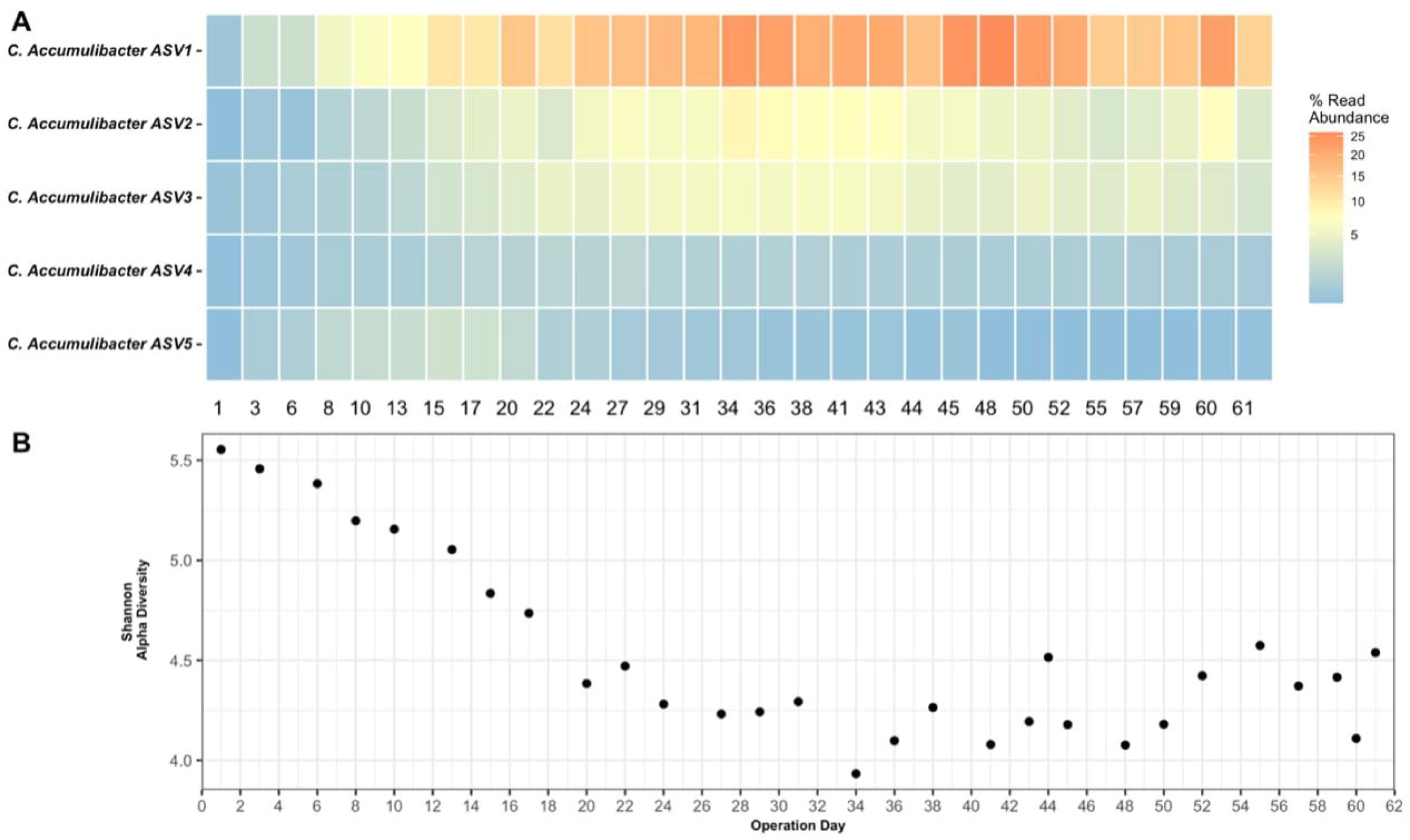
Accumulibacter ASVs and Shannon Diversity over the Time series. **A)** The top five Accumulibacter ASVs were plotted over the course of the time series. Relative abundance is plotted as the total number of reads mapping back to the ASV. **B)** Shannon’s alpha diversity calculated for each sample over the course of the time series.

**Supplementary Figure 3.**
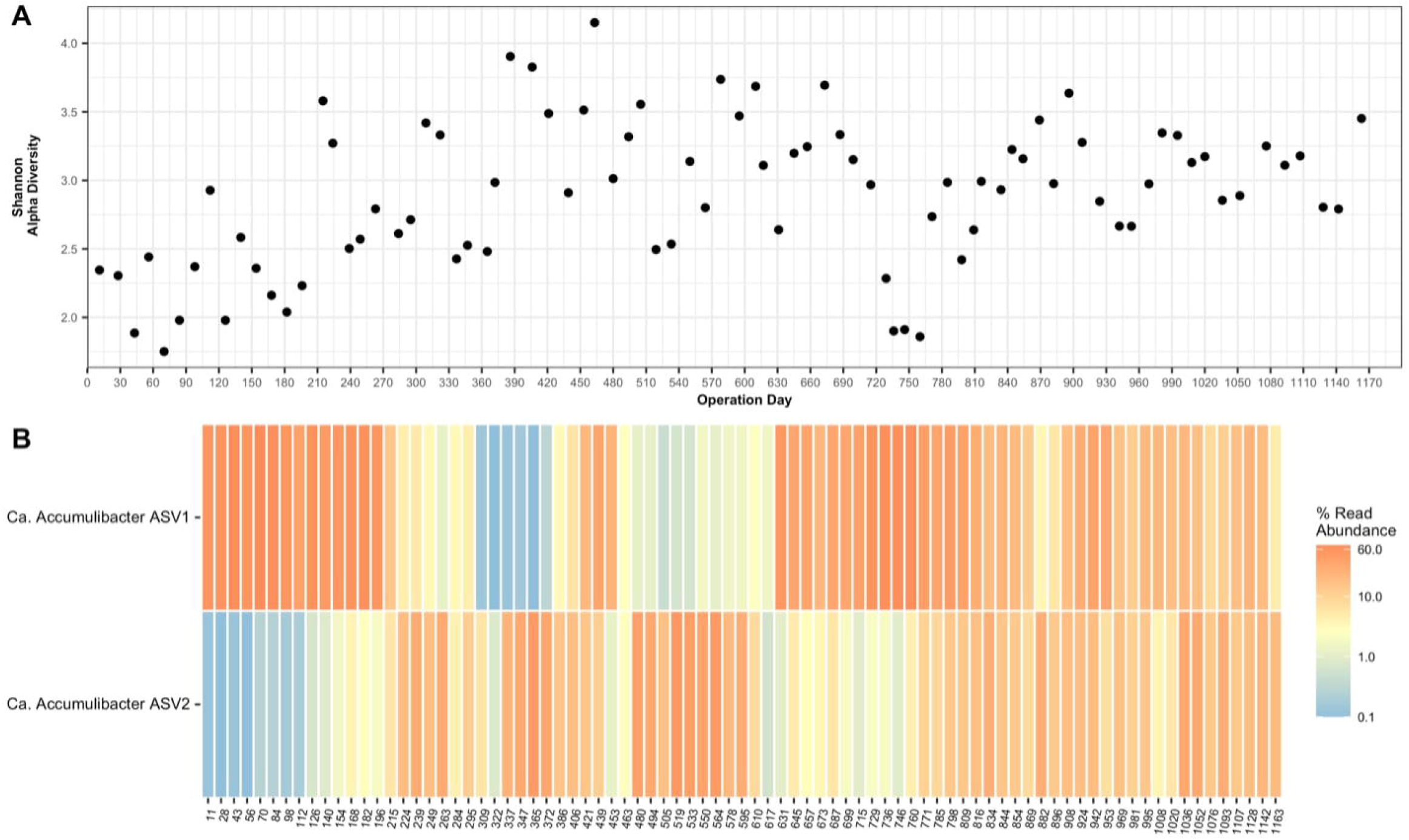
Shannon’s Diversity and Accumulibacter ASVs over the Time series. **A)** Shannon’s alpha diversity calculated for each sample over the course of the time series. **B)** Top two most abundant ASVs belonging to *Ca*. Accumulibacter and relative abundance over the time series. Read abundance refers to total reads mapping to these ASVs in that sample. Dynamics of these two ASVs roughly match the dynamics of clades IA and IIA using clade-specific *ppk1* qPCR primers in Figure 2D.

**Supplementary Figure 4.**
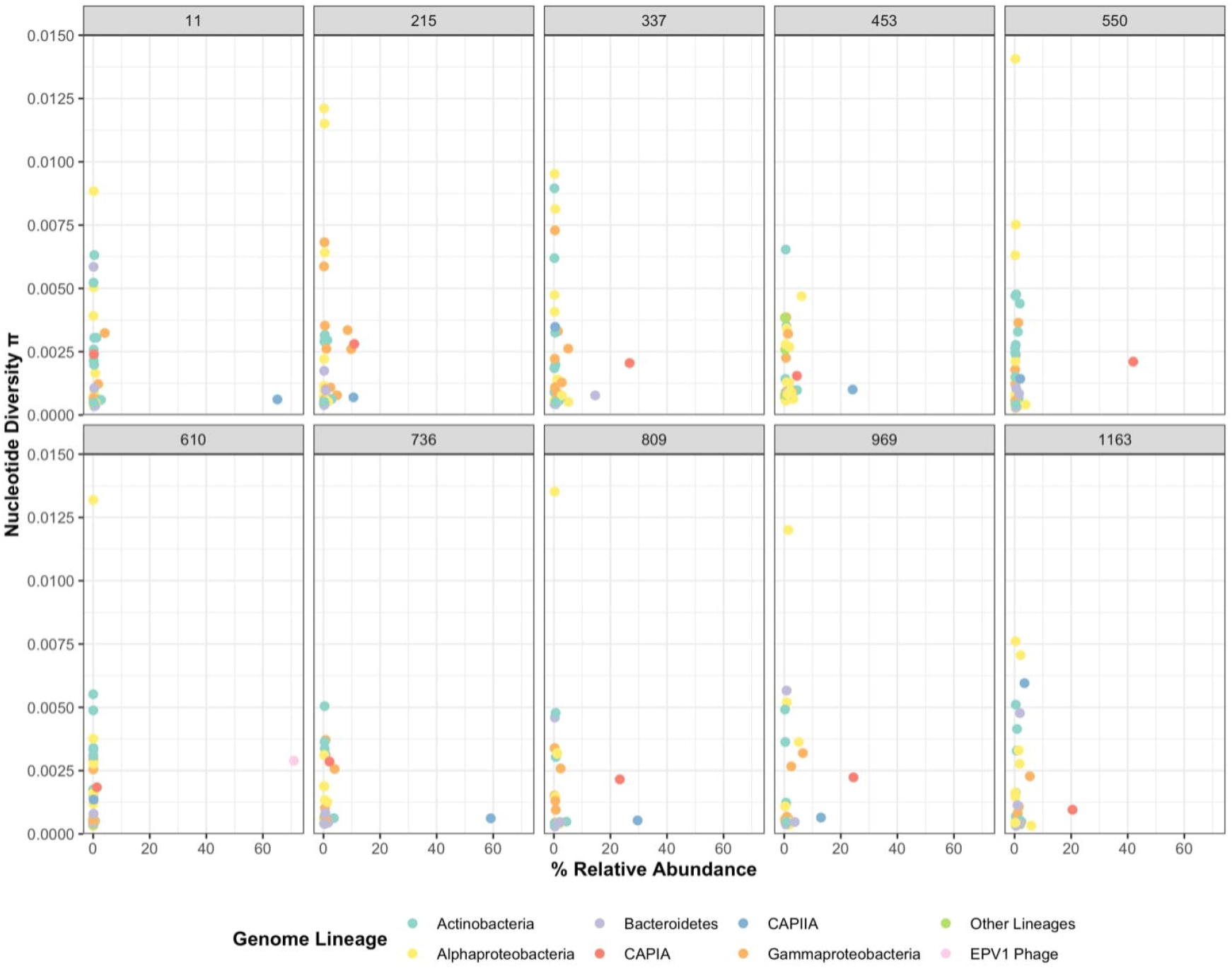
Relative Abundance and Nucleotide Diversity of MAGs in each sample. Comparisons of relative abundance and within-sample nucleotide diversity for each MAG in each sample that was above the coverage threshold of 10X coverage in that sample.

**Supplementary Figure 5.**
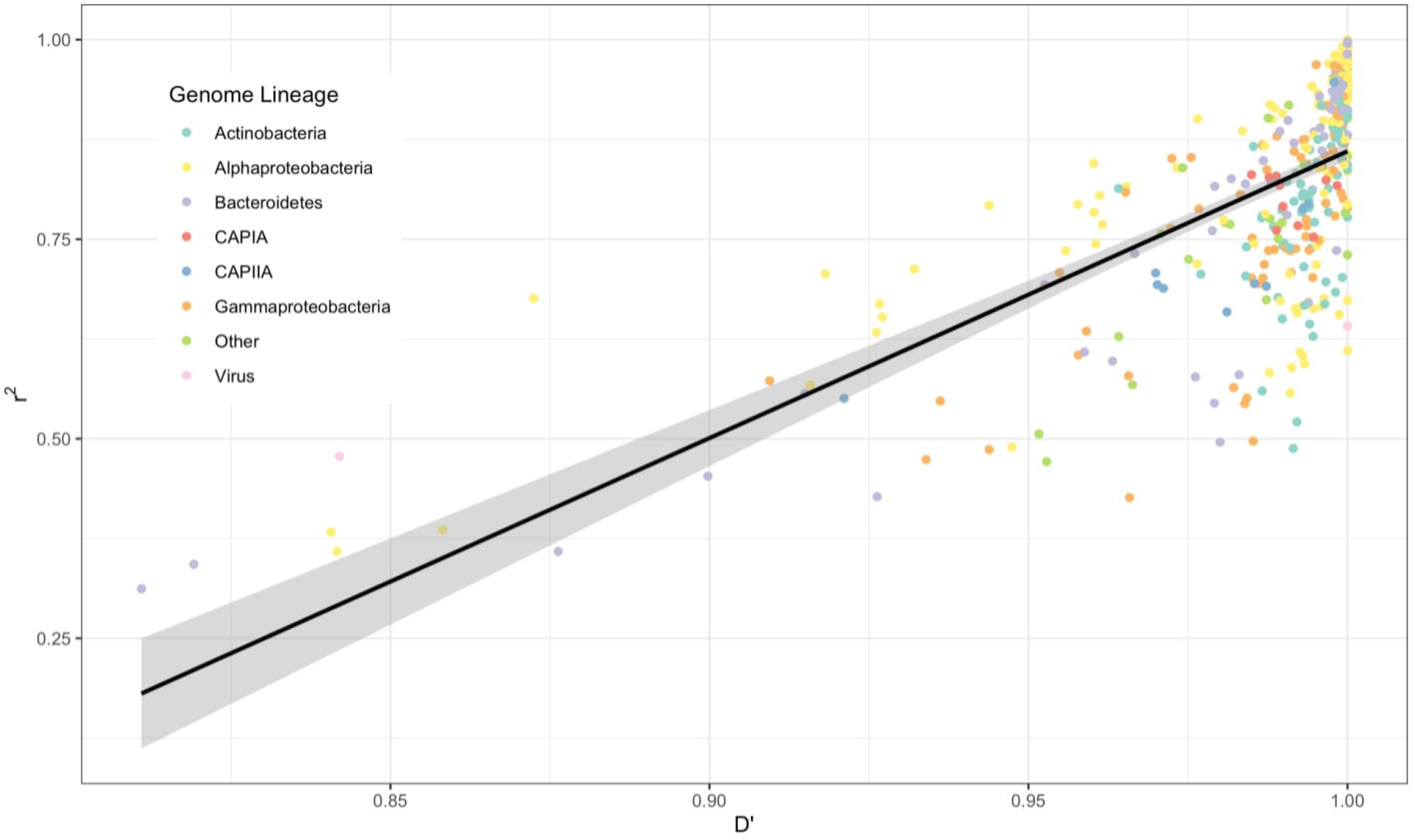
Comparison of Linkage Disequilibrium *r*^*2*^ and D’ Metrics for Interpreting Recombination. Each point represents the *r*^*2*^ and D’ value for an individual MAG in a single sample that met the coverage threshold above 10X in that sample. These two values are supposed to follow a linear relationship, where a D’ of 1 infers no recombination, and the higher the *r*^*2*^ value, the less recombination occurring in that genome. Points are colored by genome lineage matching colors in Figure 3.

